# Proof of concept study deploying CRISPR inhibition and activation opens new avenues for systematic biological exploration in zebrafish

**DOI:** 10.1101/2024.09.16.613289

**Authors:** Nelson B. Barrientos, Elyse A. Shoppell, Rachel J. Boyd, Valeria C. Culotta, Andrew S. McCallion

**Affiliations:** McKusick-Nathans Department of Genetic Medicine, Johns Hopkins University School of Medicine, Baltimore, MD 21205, USA; Department of Biochemistry and Molecular Biology, Johns Hopkins University Bloomberg School of Public Health, Baltimore, MD 21205, USA; Department of Medicine, Johns Hopkins University School of Medicine, Baltimore, MD 21205, USA

**Keywords:** Zebrafish, CRISPR-activation, CRISPR-inhibition

## Abstract

The application of CRISPR interference (CRISPRi) and CRISPR activation (CRISPRa) technologies in zebrafish has the potential to expand its capacity for the study of gene function significantly. We report proof-of-principle data evaluating transient expression of a codon optimized CRISPRi/a system for zebrafish across established pigmentary and growth phenotypes. A codon-optimized and catalytically inactive *cas9* gene (*dcas9*) was cloned upstream of codon-optimized Krüppel associated box (KRAB) and methyl-CpG binding protein 2 (MeCP2) for CRISPRi, and VP64 for CRISPRa. To validate CRISPRi, we targeted key genes in melanocyte differentiation (*sox10* and *mitfa)*; and melanin production (tyrosinase; *tyr*). Microinjection of CRISPRi mRNA and single guide RNAs (sgRNAs) targeting the *tyr* promoter or 5’-UTR resulted in larvae with hypopigmented epidermal melanocytes. Similarly, CRISPRi targeting of the *sox10 or mitfa* promoters results in hypopigmentation of epidermal melanocytes consistent with their roles upstream of *tyr,* and the role of *sox10* in activation of *mitfa*. Finally, we tested both CRISPRi/a to modulate a single gene to yield hypomorphic and hypermorphic effects, selecting *mrap2a* as our target. This gene regulates energy homeostasis and somatic growth via inhibition of the melanocortin 4 receptor gene (*mc4r*). We show that inactivating or activating *mrap2a* with CRISPRi/a significantly decreases or increases larval body length, respectively. We demonstrate that CRISPRi/a can modulate control of zebrafish gene expression, facilitating efficient assay of candidate gene function and disease relevance.

## INTRODUCTION

Zebrafish (*danio rerio*) are a powerful model system in which to study functional components of vertebrate genomes. Comparative genomics analyses have shown that 61.5% of mouse protein-coding genes and 71% of human protein-coding genes have at least one zebrafish orthologue (1,2). Additional features, including short generation time, high fecundity, external fertilization, and embryonic transparency, have distinguished zebrafish as a desirable model system in which to examine developmental processes. As such, zebrafish have been widely used to study the biological relevance of regulatory elements (3–6), and human disease-associated genes (7–14).

Emerging technologies for transgenesis and genome engineering, such as such as Gal4-UAS (15); Tol2-transposase transgenesis (3); Zinc Finger Nucleases (16); and transcription activator-like effectors (TALENs) (17) have been developed and rapidly implemented in zebrafish. Over the last decade, zebrafish have also been shown to be amenable to CRISPR-Cas-based genomic engineering applications, including targeted mutagenesis, base editing, and generation of knock-in and knockout models (18–22). More recently, the emergence of CRISPR interference (CRISPRi) and activation (CRISPRa) have presented additional opportunities to explore functional components of vertebrate genomes. CRISPRi/a involves the fusion of transcriptional effectors to a catalytically inactive, or “dead” Cas9 (dCas9). CRISPRi employs a transcriptional repressor, such as a Krüppel associated box (KRAB) domain and methyl-CpG binding protein 2 (MeCP2) (23,24), while in the case of CRISPRa, a transcriptional activator, such as VP64, is fused to the dCas9 (25). To date, the CRISPRi/a systems have seen limited application in zebrafish (26), whereas they have been more rapidly employed in mammalian organisms to target human disease-relevant genes (27–29).

Here, we report initial experiments in the development and evaluation of zebrafish codon-optimized zdCas9-KRAB-MeCP2 and zdCas9-VP64 CRISPR systems. We present a systematic evaluation of these modified proteins in functional proof-of-concept experiments by targeting two easily scored phenotypes: zebrafish pigmentation and somatic growth. In addition to providing visual confirmation of the efficacy of both CRISPRi and CRISPRa, these two model phenotypes and their corresponding pathways are well-characterized and are involved in human disease.

Specifically, we chose to test our CRISPRi system by targeting genes involved in melanocyte differentiation and pigmentation, *sox10 (SRY-*box transcription factor 10), *mitfa* (melanocyte inducing transcription factor a), and the gene for the rate-determining enzyme in pigmentation, *tyr (*tyrosinase). *sox10* directly activates the *mitfa* promoter to stimulate downstream genes involved in melanocyte differentiation (33,34); this leads to the activation of *tyr*, at approximately 16 hours post fertilization (16 hpf) to induce melanin production by 24 hpf (30–32). In the developing zebrafish embryo, due to whole-genome duplication, *mitfa* is co-expressed with *mitfb* in the retinal pigment epithelium, but only *mitfa* is expressed in neural crest cells, which differentiate to give rise to epidermal melanocytes (31,33,34).

We hypothesize that independently targeting each of these genes with functional CRISPRi will yield a visible reduction in zebrafish pigmentation. In the case of *sox10,* we expect to see marked reduction in pigmentation globally due to its role in melanocyte cell fate and activation of *mitfa* and *tyr* to promote melanin production in zebrafish and other vertebrates (35,36). Due to its role in multiple stages of melanocyte differentiation and its function as a transcription factor, silencing *mitfa* is also expected to result in phenotype of epidermal hypopigmentation (37,38). Lastly, we hypothesize that targeting the rate-determining enzyme in melanin pigment formation (*tyr)* will result in hypopigmentation of melanocytes throughout the zebrafish body (30,35,39,40). Zebrafish melanocytes have been extensively studied by our group and others (4,6,41,42), and targeting genes in this pathway is highly relevant to the study of human disease, including melanoma (43–45), Waardenburg Syndrome (46,47), microphthalmia, and Hirschsprung’s disease (48,49).

Expanding from this initial study, we sought to apply both CRISPRi and CRISPRa to *mrap2a*, a gene with known inhibitory activity to regulate somatic growth in the zebrafish embryo (50). *mrap2a* encodes the melanocortin receptor accessory protein 2a, which has been found to inhibit the transcription of the *mc4r* (melanocortin-4 receptor) (50). We hypothesize that targeting the *mrap2a* promoter with CRISPRa will increase *mrap2a* expression and therefore amplify the inhibition of *mc4r* and promote visible overgrowth phenotypes in zebrafish larvae. This is supported by evidence in human patients, wherein autosomal dominant loss of function mutations in *MCR4* results in severe childhood-onset obesity (51,52). By contrast, we hypothesize that effective CRISPRi targeting of *mrap2a* will result in decreased *mc4r* inhibition and result in undergrown zebrafish larvae, consistent with reported morpholino-based knock-down experiments (50).

Ultimately, we outline the development of zebrafish-optimized zdCas9-KRAB-MeCP2 and zdCas9-VP64 CRISPR constructs. We establish their utility for efficient screening of candidate gene function and disease relevance. Ongoing work in the lab will establish stable transgenic lines expressing these proteins to facilitate rapid and systematic screening of gain and loss of gene function by leveraging custom guide RNAs (gRNAs) to target promoters, 5’-UTR, or cell-dependent cis-regulatory elements of selected candidate genes.

## RESULTS

### CRISPRi modulates tyrosinase expression and its effect is observable between the prim 20-25 stages

To determine the functional capacity of zebrafish codon-optimized CRISPRi (Fig. 1A), one-cell stage embryos of wild-type AB (WT AB) zebrafish were injected with CRISPRi mRNA and target sgRNAs. As part of a series of proof-of-concept experiments, we designed sgRNAs targeting the promoter region and 5’-UTR of the tyrosinase (*tyr*) gene, predicting that inhibition would lead to diminished larval pigmentation, similar to established loss of function studies at this locus (30,32,40).

**Figure 1:**
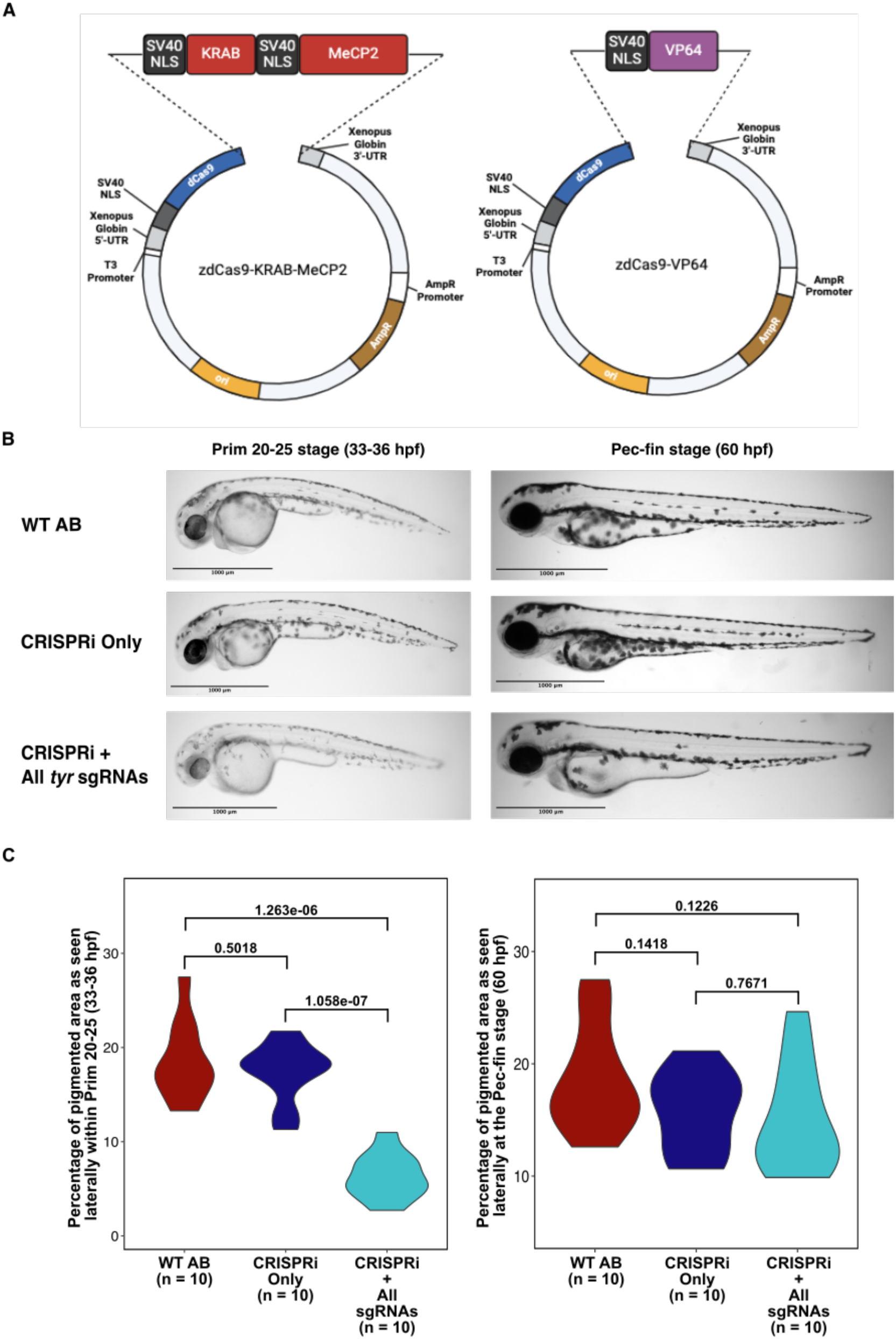
CRISPRi/a construct design and *tyr* silencing. **A.** Zebrafish codon-optimized plasmids were synthesized using NEB HiFi DNA assembly, transformed and amplified in NEB 5-α chemically competent cells. Plasmid DNA was linearized and *in vitro* transcribed before microinjection into one-cell embryos. **B.** Zebrafish embryos injected with CRISPRi (final concentration 100-300 ng/µl) + cocktail of sgRNAs (final concentration 150 ng/µl) targeting *tyr* assayed at prim 20-25 (33-36 hpf) and pec-fin (60 hpf) stages for differences in pigmentation patterns. The larvae injected with CRISPRi and sgRNAs displayed reduced pigmentation at the prim 20-25 stage when compared to controls. The effect of CRISPRi dissipated by the pec-fin stage as observable pigmentation patterns were indistinguishable. Scale bars 1000 µm. **C.** Pigment analysis of larvae at Prim 20-25 stage (33-36 hpf) and Pec-fin stage (60 hpf) to measure the percentage of pigmented area, as viewed laterally. At Prim 20-25, the percentage of pigmented area is statistically significant (Welch two-sample t-test); by contrast, at Pec-fin, while not statistically significant, CRISPRi+sgRNA injected embryos show less percentage of pigmented area than WT AB and injection controls.

Initially, sgRNAs were designed to target the *tyr* promoter or 5’-UTR (Methods) and one-cell embryos (n ≥ 200) were injected with CRISPRi (100-300 ng/µl, final concentration) and a cocktail of all five sgRNAs (150 ng/µl/each, final concentration). Zebrafish larvae assayed at the prim 20-25 stages (33-36 hpf) displayed marked reduction in pigmentation of melanocytes (Fig. 1B), consistent with phenotypes arising from loss of function alleles in zebrafish (30) and other vertebrates (39). By contrast, larvae receiving only the CRISPRi mRNA (n ≥ 200) or uninjected (WT AB) controls (n ≥ 200), displayed no diminution in melanization at these stages. As expected, the observed suppression of *tyr* dissipated over time, and by the pec-fin stage (60 hpf), reduction in melanization was less noticeable in larvae receiving CRISPRi plus the cocktail of all sgRNAs (Fig. 1B).

We measured the percentage of pigmentated area, imaged laterally, from a sample (n = 10) of larvae from each condition, at the prim 20-25 stage, and found a statistically significant reduction of pigmentation between uninjected control (WT AB) and larvae injected with CRISPRi plus all *tyr* sgRNAs (*p-value* = 1.263e-06, Fig. 1C, Supplemental Table 4a). We also observed a statistically significant reduction of pigmentation between injected control (embryos injected with CRISPRi mRNA only) and larvae injected with CRISPRi plus all *tyr* sgRNAs (*p-value* = 1.058e-07, Fig. 1C, Supplemental Table 4a). We did not observe a statistically significant difference between injected and uninjected controls (*p-value* = 0.5018, Fig. 1C, Supplemental Table 4a). We subsequently performed the same pigmentation analysis at the pec-fin stage and observed the same trend of hypopigmentation in CRISPRi plus all *tyr* sgRNAs (Fig. 1B-C, Supplemental Table 4b). However, the results were not statistically significant across conditions (*p-values* = 0.1226, 0.1418, and 0.7671, respectively, Fig. 1C, Supplemental Table 4b), confirming that the effects of CRISPRi mRNA injection diminish over time.

We tested if delivery of each individual *tyr* sgRNAs could similarly silence transcription of *tyr* and reduce melanin production. One of the five sgRNAs that we tested, sgRNA4 targeting the promoter, efficiently modulated *tyr* expression (Supplemental Fig. 1), resulting in hypopigmentation within prim 20-25 (33-36 hpf). We used this sgRNA to assess the optimal concentration of CRISPRi and injected larvae each with 100 ng/µl, 300 ng/µl, and 500 ng/µl of CRISPRi along with sgRNA4 (150 ng/µl). Our observations suggest no advantage to injecting more than 300 ng/µl (final concentration) of CRISPRi mRNA (Supplemental Fig. 1). These results demonstrate the effectiveness in modulating *tyr* by employing our zebrafish codon-optimized CRISPRi with individual or pooled sgRNAs to target a gene’s promoter. To build on these observations, we sought to confirm the hypopigmentation phenotype by targeting other key pigment genes that act upstream and are known to regulate *tyr*.

### CRISPRi targeting of *sox10* impedes epidermal pigmentation zebrafish

Having established that *tyr* can be modulated via CRISPRi, we sought to evaluate a broader applicability of our strategy, evaluating the effect of modulating genes upstream from *tyr* in melanocyte differentiation, for which we predict overlapping phenotypic outcomes. We first investigated whether *sox10*, the gene that encodes for a transcription factor known to be key to promote melanin production in zebrafish and other vertebrates (35,36) could be silenced via CRISPRi. We designed sgRNAs to target the promoter of the *sox10* gene, predicting that silencing it would lead to downregulation of downstream pigment genes, *mitfa* and *tyr*, and subsequent hypopigmentation in the zebrafish.

Four sgRNAs were designed to target the *sox10* promoter (Methods). Zebrafish larvae (n ≥ 200) injected with CRISPRi (100-300 ng/µl, final concentration) plus a cocktail of all *sox10* sgRNAs (150 ng/µl/each, final concentration) displayed marked reduction in pigmentation of epidermal melanocytes, assayed at prim 20-25 (33-36 hpf), consistent with reduction of pigmentation when targeting downstream gene *tyr* (Fig. 2A, Supplemental Fig. 2). Furthermore, we observed that sgRNA3 was effective at reducing pigmentation when injected individually with CRISPRi (Fig. 2A, Supplemental Fig. 2). We measured the percentage of pigmented area, imaged laterally at the prim 20-25 stage, from a sample (n = 10) of larvae from each condition and found a statistically significant reduction of pigmentation between uninjected control (WT AB) and larvae injected with CRISPRi plus all *sox10* sgRNAs and CRISPRi plus *sox10* sgRNA3 (*p-value* = 7.389e-07, 2.724e-06, respectively, Fig. 2B, Supplemental Table 5a). We also observed a statistically significant reduction of pigmentation between injected control (embryos injected with CRISPRi mRNA only) and larvae injected with CRISPRi plus all *sox10* sgRNAs and CRISPRi plus *sox10* sgRNA3 (*p-value* = 8.704e-09, 3.243e-06, respectively, Fig. 2B, Supplemental Table 5a). Further, we did not observe a statistically significant reduction in pigmentation between larvae receiving only the CRISPRi mRNA (n ≥ 200) and uninjected (WT AB) controls (n ≥ 200) (*p-value =* 0.097, Fig. 2B, Supplemental Table 5a). The effects of CRISPRi dissipated by the pec-fin stage (60 hpf) as the pigmentation patterns between injected and uninjected were indistinguishable (Fig. 2A, Supplemental Table 5b). When measuring the percentage of pigmented area at this stage, we only observed at statistically significant reduction in pigmentation between WT AB and CRISPRi plus all *sox10* sgRNAs (*p-value =* 0.019, Fig. 2B, Supplemental Table 5b). We did not observe a statistically significant reduction in pigmentation across the other conditions (*p-value* = 0.481, 0.090, 0.999, 0.454, and 0.102, respectively, Fig. 2B, Supplemental Table 5b), which confirms our observation that as larvae grow, the effects of CRISPRi mRNA injections dissipate. It is noteworthy to mention that larvae injected with CRISPRi plus sgRNAs appear to be slightly developmentally delayed as compared to controls, which may be the result of injection variability, nucleic acid toxicity, or off-target effects. This is also consistent with an observed reduction in retinal pigment epithelium (RPE) pigmentation within Prim 20-25 in larvae injected with CRISPRi plus sgRNAs as well as in those injected with CRISPRi alone, as compared to WT (*p-value* = 0.048, Supplemental Fig. 2B, Supplemental Table 6). Given that these effects are not consistent with the established loss of function phenotypes for these genes (33) , we do not score or interpret RPE phenotypes throughout. However, the overall results are in line with previous work that demonstrated that *sox10* is important for melanocyte development and melanin production with its role in activating *tyr*, the rate-determining enzyme in melanin production (30,35,39,40). To exclude the possibility of off-target effects by *sox10* sgRNA, we employed sgRNA3 and standard CRISPR/Cas9 to establish correct targeting of the *sox10* promoter. Employing a standard T7 endonuclease I (T7EI) assay (Methods) we confirm that sgRNA3 is effective at guiding Cas9 to the *sox10* promoter to induce double-strand breaks. We collected various clutches (n = 50) of WT AB, CRISPR/Cas9 only, and CRISPR/Cas9 plus sgRNA3 and extracted genomic DNA to amplify the *sox10* promoter. PCR amplification (Oligonucleotide primers: Forward: 5’-CTCCCAACTCTTATTAACGA-3’; Reverse: 5’-ACACAGAGTAAGCGATAGAG-3’) of this locus and subsequent treatment with T7EI resulted in heteroduplex formation, indicating effective targeting of *sox10* (Supplemental Fig. 2C).

**Figure 2:**
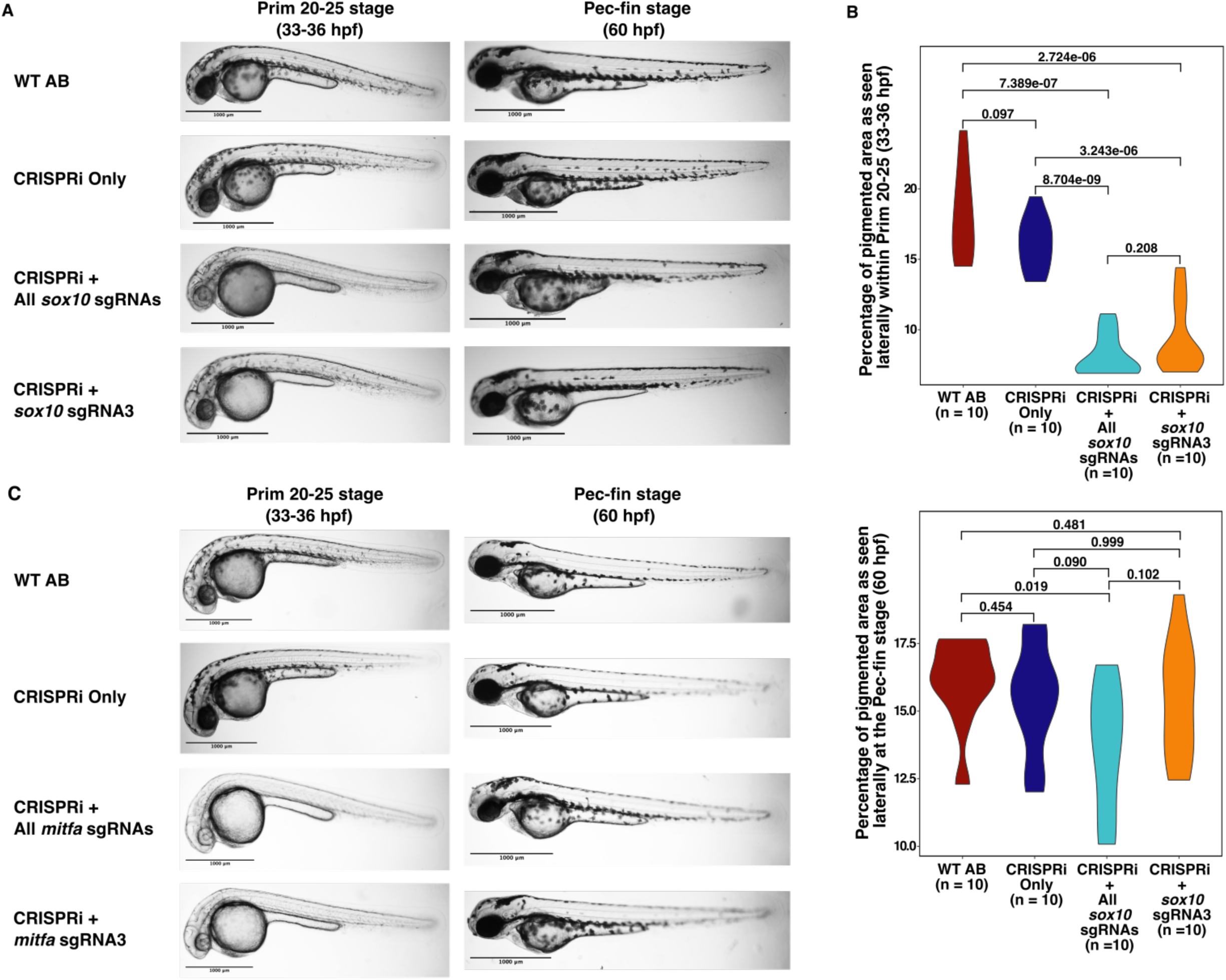
CRISPRi targeting the promoters of the *sox10* and *mitfa* genes. **A.** Zebrafish 1-cell embryos injected with CRISPRi + sgRNAs targeting *sox10* assayed at prim 20-25 (33-36 hpf) and pec-fin (60 hpf) stages for differences in pigmentation patterns. WT AB and injected controls did not show reduced pigmentation around the prim 20-25 stages, whereas embryos injected with CRISPRi + All *sox10* sgRNAs or CRISPRi + *sox10* sgRNA3 were hypopigmented. **B.** At the prim 20-25 stage, differences in pigmentation patterns across conditions are statistically significant (*p-value* < 0.05). At the pec-fin stage, the effect of CRISPRi dissipated as patterns of hypopigmentation are less variable with the only statistically significant difference being between WT AB and CRISPRi + All *sox10* sgRNAs based on Welch two-sample t-test. **C.** Zebrafish embryos injected with CRISPRi + sgRNAs targeting the *mitfa* promoter. Larvae were assayed at prim 20-25 and pec-fin stages for differences in pigmentation patterns. At this stage, embryos injected with CRISPRi + All sgRNAs or CRISPRi + sgRNA3 displayed a hypopigmented pattern as compared to injected and uninjected controls. At the pec-fin stage, the effect of CRISPRi has dissipated as patterns of hypopigmentation displayed no significant differences across conditions. Scale bars: 1000 µm.

### Effective CRISPRi modulation of *mitfa* limits epidermal melanin production

Extending from our results modulating *sox10* and *tyr* expression, we next evaluated whether targeting the gene encoding the transcription factor mitfa would also induce hypopigmentation. The *mitfa* gene plays critical roles in pigmentation throughout vertebrates (*MITF/Mitf*), activating *tyr* transcription during early development (35,59). We designed sgRNAs to target the promoter and 5’-UTR regions of the *mitfa* gene, predicting that, as with known loss-of-function (LoF) alleles (34,59), titrating activity of this transcription factor would also result in reduced activation of *tyr* in epidermal melanocytes and thus lower melanization across the body.

Four sgRNAs were designed to target the *mitfa* promoter (Methods). As before, zebrafish one-cell embryos (n ≥ 200) injected with CRISPRi (100-300 ng/µl, final concentration) and a cocktail of all *mitfa* sgRNAs (150 ng/µl/each, final concentration) displayed marked reduction in pigmentation of epidermal melanocytes, assayed at prim 20-25 (33-36 hpf) (Fig. 2C). This is consistent with prior *mitfa* LoF studies (60) where failure to express *mitfa* prevents the production of melanocyte precursors, leading to reduced global pigmentation in zebrafish larvae. In contrast, no reduction in pigmentation was observed in larvae receiving only the CRISPRi mRNA (n ≥ 200) or uninjected (WT AB) controls (n ≥ 200) (Fig. 2C). Additionally, a single *mitfa* sgRNA targeting the promoter or 5’-UTR effectively silenced *mitfa,* resulting in marked reduction of melanin production at prim 20-25 (33-36 hpf) (Supplemental Fig. 3). Silencing of *mitfa* dissipated over time and by the pec-fin stage (60 hpf), melanization was similar between controls and injected larvae (Fig. 2C, Supplemental Fig. 3). These results are consistent with the hypopigmentation phenotype in epidermal melanocytes that is observed when modulating the *tyr* gene (Fig. 1B-C) and transcription of *sox10* (Fig. 2A-B).

### CRISPRi/a affects zebrafish length by modulating transcription of *mrap2a*

To build upon the successful demonstration of CRISPRi-mediated modulation of zebrafish pigmentation, we set out to test our CRISPRa (Fig. 1A) system focusing on somatic size (length). The benefit of this phenotype is that it provides the opportunity for evaluation of both reduction and increase in gene function. We targeted the *mrap2a* gene which is known to regulate the expression of the melanocortin 4 receptor (*mc4r*) gene by downregulating its transcription (50,62). Both genes have been shown to be involved in energy homeostasis and somatic growth in mice and zebrafish; and are thought to be involved in a significant portion of familial early-onset obesity in humans (63). We predicted that targeting *mrap2a* with CRISPRa and further increasing its transcription would result in a decrease in *mc4r* transcription, which would lead to an increase in larval length. By contrast, downregulating *mrap2a* with CRISPRi would lead to an increase in *mc4r* expression, resulting in larvae shorter in length (50,63).

While a range of loss-of-function experiments have been performed targeting *mrap2a* (50,62,63), gain-of-function (GoF) experiments targeting *mrap2a* have not been explored. We designed four sgRNAs to target the *mrap2a* promoter (Methods) with our CRISPRa system. Zebrafish larvae (n ≥ 200) injected with CRISPRa (100-300 ng/µl, final concentration) and a cocktail of all *mrap2a* sgRNAs (150 ng/µl/each, final concentration) displayed marked increase in body length, assayed at the pec-fin stage (60 hpf) (Fig. 3A). Similarly, we tested if an individual sgRNA could guide the dCas9-VP64 to modulate body length and found sgRNA3 capable of inducing an observable phenotype (Fig. 3A-B). By contrast, no increase in larval length was observed at the pec-fin stage (60 hpf) in larvae receiving only the CRISPRa mRNA (n ≥ 200) or in uninjected (WT AB) controls (n ≥ 200). Moreover, the effects of CRISPRa were still noticeable at the day 5 stage (120 hpf) and at the day 7 stage (168 hpf), as the length of these larvae was still longer than control conditions (Fig. 3A, Supplemental Fig. 4). Furthermore, we measured the length of larvae receiving CRISPRa plus sgRNA3 (n = 50), sgRNA3-only (n = 50), and uninjected (WT AB) controls (n = 50) and performed a Welch two-sample t-test to analyze if the average length between these conditions was statistically different (Fig. 3B, Supplemental Table 7). At the pec-fin stage (60 hpf), the average length for these conditions was 3673.67 μm, 3468.86 μm, and 3466.28 μm (*p-value* = 8.58e-16; 1.83e-14, and 0.9181), respectively. The longer larval length phenotype due to increased *mrap2a* transcription and decreased *mc4r* expression is consistent with increased size and weight in mice that become obese due to *mc4r* haploinsufficiency (29), and with morbid obesity cases in humans due to frameshift (loss of function) mutations in *mc4r* (64).

**Figure 3:**
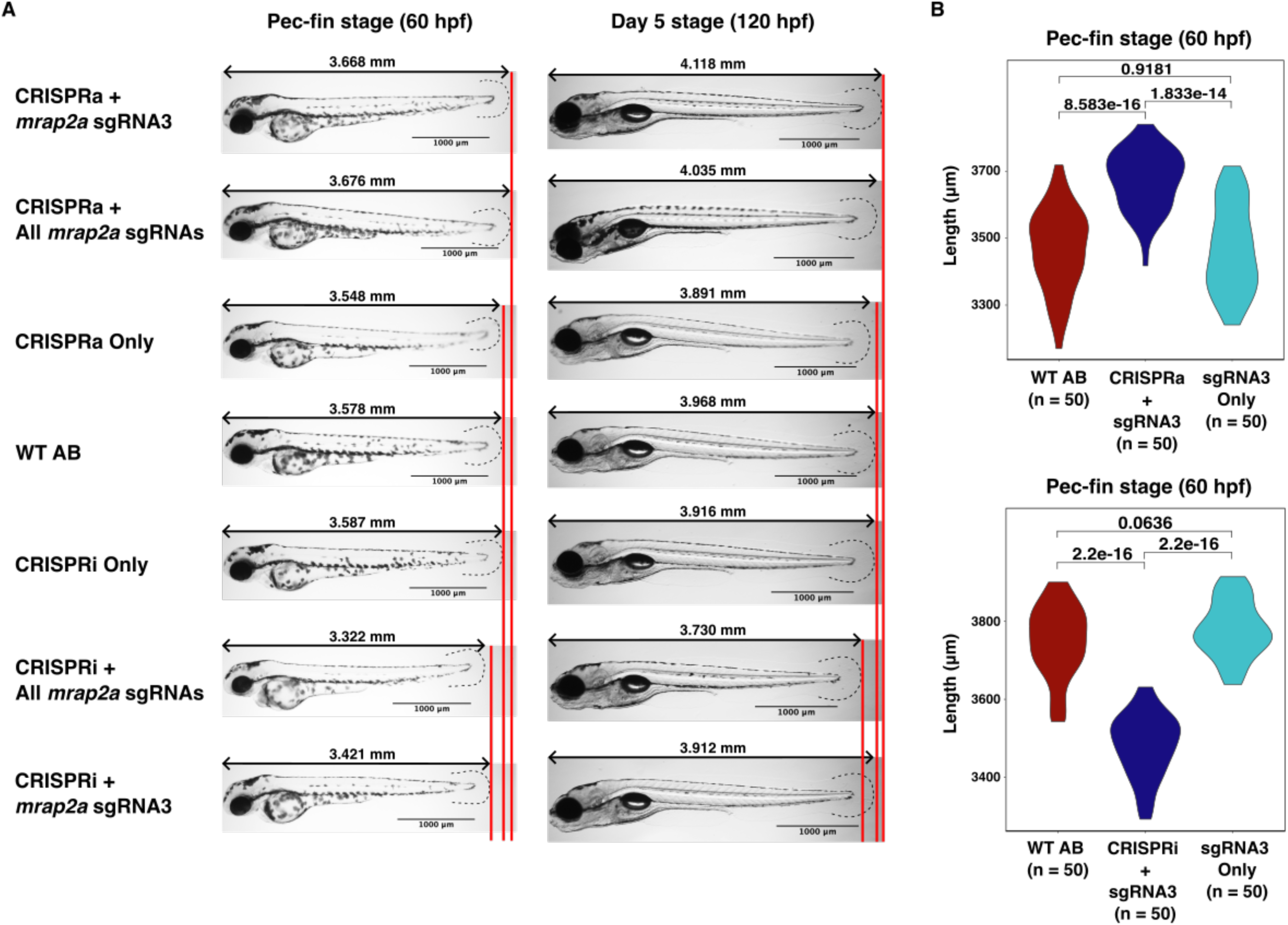
Targeting the promoter of *mrap2a* with CRISPRa and CRISPRi. **A.** Length comparison between the pec-fin and day-5 stages between WT AB embryos and embryos injected with CRISPRa and CRISPRi with all *mrap2a* sgRNAs or only sgRNA3. Length measured laterally from tip of the mouth to end of translucent tail fin (dashed lines). Red lines depict length differences between controls and experimental conditions. Scale bars: 1000 µm. **B.** At the pec-fin stage, 50 larvae per condition were measured in length from the tip of the mouth to the end of the translucent tail fin. The Welch two-sample t-test was used to compare the average length between conditions.

We used the same sgRNAs to target and impede *mrap2a* and increase *mc4r* expression. Zebrafish larvae (n ≥ 200) injected with CRISPRi (100-300 ng/µl, final concentration) plus all sgRNAs (150 ng/µl/each, final concentration) displayed a decrease in larval length at the pec-fin stage (60 hpf) (Fig. 3A, 3C), which is consistent with rescuing obese mice with *mc4r* haploinsufficiency (29). Additionally, sgRNA3 was able to guide the CRISPRi to modulate silencing of *mrap2a* individually and led to a decrease in larval length, observed at the pec-fin stage (60 hpf) (Fig. 3A. 3C). By contrast, larval length observed at the pec-fin stage (60 hpf) for larvae receiving only the CRISPRi mRNA (n ≥ 200) or in uninjected (WT AB) controls (n ≥ 200) was higher than our experimental condition (Fig. 3A). Further, larvae injected with CRISPRi still displayed a shorter length around the day 5 stage (120 hpf); however, their length was similar to controls at the day 7 stage (168 hpf) (Fig. 3A, Supp Fig. 4). We also measured the larval length for the conditions CRISPRi plus sgRNA3 (n = 50), sgRNA3-only (n = 50), and uninjected (WT AB) controls (n = 50) at the pec-fin stage (60 hpf) and found their average length to be 3481.98 μm, 3779.96 μm, and 3748.82 μm, respectively (Fig. 3B, Supplemental Table 8). We analyzed the statistical significance between these means with a Welch two-sample t-test and found the difference in means between experimental and control conditions to be statistically significant (*p-value* = 2.2e-16, 2.2e-16, and 0.0636). Although highly significant, the observed magnitude of effect may potentially be further enhanced by simultaneous modulation of the agouti-related protein (agrp), which Sebag et al. demonstrated acted in concert with *mrap2a* in antagonism of *mc4r* (50).

## DISCUSSION

CRISPRi/a has emerged as a powerful tool in genetic research, offering the ability to modulate gene expression both for exploration of fundamental mechanisms and for potential therapeutics. It offers the capacity to modulate gene expression without inducing genetic alterations which makes CRISPRi/a valuable technology for the study of gene regulation and for efficient exploration of both gain and loss of function phenotypes. Nevertheless, their development and application in zebrafish have been slow over the past decade, with only one study describing an effort to implement it (26), while many others apply CRISPR-based editing technologies (18,19,21,53,54,65).

In this study, we chose to fuse a catalytically inactive Cas9 with the transcriptional repressor KRAB-MeCP2. Our choice to use dCas9-KRAB-MeCP2 was guided by previous studies that demonstrated its ability to induce gene silencing while outperforming dCas9-KRAB systems (24,66) implemented in previous CRISPRi experiments in zebrafish (26). In contrast, for our CRISPRa system, we fused the dCas9 with the transcriptional activator VP64. Comparisons between different Cas9 activators have demonstrated the VP64 to effectively induce gene activation (25). While other Cas9 activators such as VPR, SAM, and Suntag have been shown to outperform VP64, these systems have not been tested in zebrafish and/or require special sgRNAs to induce their effect. By using a catalytically inactive Cas9 fused with the transcriptional repressors KRAB-MeCP2 or the transcriptional activator VP64, we have achieved transcriptional control over key genes involved in melanocyte differentiation (*sox10* and *mitfa*), pigmentation (*tyr*), and somatic growth (*mrap2a*) in zebrafish. Our results underscore the versatility and efficiency of CRISPRi/a in zebrafish research. For instance, by independently downregulating *sox10, mitfa,* and *tyr*, we established zebrafish larvae with marked reduction in pigmentation. Similarly, by activating or downregulating *mrap2a*, we successfully modulated larval length.

These proof-of-principle experiments underscore the potential of CRISPRi/a technology to control gene expression by targeting gene promoters. CRISPRi/a is relatively straightforward to implement compared to other genetic editing technologies, which further increases its appeal. In addition, implementing this technology in transparent one-cell zebrafish embryos makes it easier to study the effects of gene silencing and activation during development. Further, CRISPRi/a can be implemented to study the effects of silencing or activating noncoding regulatory sequences such as enhancers, repressors, and insulators. Taking advantage of the fact that approximately 70% of human genes have a zebrafish gene homolog facilitates the study of gene regulatory mechanisms and networks in yet another vertebrate system. The versatility to study gene expression modulation, by either targeting gene promoters or any potential noncoding regulatory sequences in a vertebrate system, can offer insights into disease pathogenesis and therapeutic targets that may translate to clinical applications. Further, our evaluation of *mrap2a* modulation also demonstrates that modulating the regulation of one gene can elicit indirect regulation of other target genes, opening the door to a better understanding of gene regulatory networks and their exploration for potential therapeutic interventions.

CRISPRi/a technologies, just like CRISPR gene editing, can also be limited by off-target effects and variability in efficiency across genomic loci. Additionally, the ethical implications of gene expression modulation and the potential for unintended consequences underscore the importance of careful consideration in the application of this technology. In this current study, the primary limitation is the delivery of CRISPRi/a via mRNA microinjections, which is titrated as the embryo develops and the larvae grow older. Furthermore, developmental delays can occur in larvae injected with foreign mRNA, which further reflects the variability induced by the combination of mRNA microinjections, potential off-target effects, and nucleic acid toxicity. This results in a diminution of molecular and observable phenotypes and hinders the potential to study long-term effects of gene expression modulation, making it difficult to elucidate any potential disease relevance. To this end, we are actively generating stable transgenic lines expressing CRISPRi/a to facilitate rapid and systematic screening of gain or loss of gene function. Our ongoing work will expand the zebrafish toolkit, establishing the utility of CRISPRi/a for evaluation of candidate gene function and disease relevance.

## METHODS

### Generation of catalytically inactive Cas9 plasmid vector

The zebrafish optimized *Cas9* gene within the plasmid vector pT3TS-nCas9n (Addgene #46757) was mutated to produce a catalytically inactive dead *Cas9* (dCas9). The DNA sequence corresponding to the RuvC-like and HNH nuclease domains of the Cas9 protein were modified by inducing a change from Aspartic acid to Alanine at position 10 (D10A), and a change from Histidine to Alanine at position 839 (H839A). This was accomplished by implementing the NEBuilder HiFi DNA Assembly (Cat. # E5520S) protocol using oligonucleotides primers (Supplemental Table 1) generated using the NEBuilder Assembly Tool (v.2.10.1). The modified plasmid vector was then transformed into NEB 5-a competent *E. coli* cells (Cat. # C2987I) and later mini-prepped using NEB Monarch Plasmid Miniprep Kit (Cat. # T1010L). The plasmid vector generated was named pT3TS-ndCas9n.

### Codon Optimization of KRAB-MeCP2 and VP64 domains for zebrafish usage

The DNA sequence for the KRAB-MeCP2 repressor was obtained from the dCas9-KRAB-MeCP2 plasmid vector (Addgene #110821); while the DNA sequence for the VP64 activator was obtained from the dCas9-VP64-GFP plasmid vector (Addgene #61422). The codon optimization tool from IDT was used to optimize these sequences for zebrafish. The KRAB-MeCP2 repressor sequence (1,263 bp) containing an SV40 NLS before and after KRAB was optimized and ordered as a gBlocks HiFi Gene Fragment. The VP64 activator sequence (225 bp) containing an SV40 NLS before VP64 was optimized and ordered as a gBlocks Gene Fragment.

### CRISPRi/a plasmid construct generation and *in vitro* transcription

The codon optimized KRAB-MeCP2 repressor and VP64 activator were cloned into the pT3TS-ndCas9n plasmid vector containing a catalytically inactive dCas9 sequence. Cloning was carried out by employing NEBuilder HiFi DNA Assembly (Cat. # E5520S) using oligonucleotide primers (Supplemental Table 2) that overlapped both the target region of the plasmid vector and the optimized sequences. Once the optimized sequences had been cloned in, the plasmid constructs were transformed into NEB 5-a competent *E. coli* cells (Cat. # C2987I) and later mini-prepped using NEB Monarch Plasmid Miniprep Kit (Cat. # T1010L). The plasmid constructs generated were named zdCas9-KRAB-MeCP2 and zdCas9-VP64 for CRISPRi and CRISPRa, respectively.

The restriction enzyme XbaI (NEB Cat # R0145) was used to linearize the CRISPRi/a plasmid constructs. Subsequently, *in vitro* transcription of the d*Cas9* gene plus the repressor or activator was performed by employing the mMessage mMachine T3 Transcription kit (Cat. # AM1348). RNA was recovered by using the RNA Clean-up protocol from the Qiagen RNeasy Mini Kit (Cat. # 74104) and stored at -80°C.

### Designing targeted gRNAs for CRISPRi/a

Multiple single guide RNAs (sgRNAs) were designed to target the promoter or 5’-UTR of the various genes in the zebrafish. All sgRNAs followed a core 5’-N18-3’(53) sequence and were designed using the online tool CRISPOR (PMID: 29762716). At the 5’ end of the forward oligonucleotide, the nucleotides 5’-TAGG-3’ were added. At the 5’ end of the reverse oligonucleotide (reverse complement of N18 sequence), the nucleotides 5’-AAAC-3’ were added. The final form of the oligonucleotides was 5’-TAGG-N18-3’ for the forward oligo; and 5’-AAAC-N18RevCompl-3’. The oligo primers for each sgRNA were ordered from IDT (Supplemental Table 3). Oligonucleotide primers were first annealed and subsequently cloned into the pT7-gRNA plasmid vector (Addgene #46759) containing a gRNA scaffold, according to restriction enzyme cloning protocol specifications from previous studies(54). After cloning, each plasmid construct generated for each sgRNA was transformed into NEB 5-a competent *E. coli* cells (Cat. # C2987I) and later mini-prepped using NEB Monarch Plasmid Miniprep Kit (Cat. # T1010L). Linearization of these constructs was performed using the restriction enzyme BamHI-HF (NEB Cat # R3136). The linear DNA was purified using the Zymo Research DNA Clean & Concentrator-5 protocol (Cat. # D4013). Subsequently, *in vitro* transcription of linear DNA was performed using the Thermo Fisher MEGAshortscript T7 Kit (Cat. # AM1354) or the NEB HiScribe T7 High Yield RNA Synthesis Kit (Cat. #E2040S). After in vitro transcription, RNA was recovered by using the RNA Clean-up protocol from the Qiagen RNeasy Mini Kit (Cat. # 74104) and stored at -80°C.

### Zebrafish husbandry and microinjections of one-cell stage embryos

We used the zebrafish AB strain (ZFIN ID: ZDB-GENO-960809-7) as our wild-type fish (WT AB). The day before injections, adult AB zebrafish crosses were set up in tanks with dividers at a ratio of 2:1 or 3:2 females per male. The morning of injections and when the lights came on, the dividers were removed to allow for mating among zebrafish. Zebrafish embryos were injected during the first 15-20 mins after being collected to ensure injection occurred at the one-cell stage. The mixture injected to the embryos contain CRISPRi/a mRNA (100-300 ng/µl, final concentration), sgRNA mRNA (150 ng/µl/each, final concentration), 0.2 µl of Phenol Red, and Nuclease Free Water up to 5 µl. After microinjections, zebrafish embryos were incubated at 28°C in E3 embryo medium provided by Johns Hopkins FinZ Zebrafish Core Center.

### Zebrafish length measurement and image collection

All zebrafish larvae used for measurements or image collection were staged according to anatomical hallmarks described in Kimmel *et al*(55). Larvae were anesthetized using ethyl-3-aminobenzoate methanesulfonate salt (Tricane/MS-222; Sigma Aldrich #E10521) prepared at a concentration of 4 mg/mL using 1 M Tris (pH 9, 4% v/v). The Nikon AZ100 Multizoom fluorescent microscope was used for both measurements and image collection at a ratio of 3.22 μm/px. For all images, zebrafish larvae were suspended in 3% Methyl Cellulose (Sigma Aldrich #M0387) solution. Larvae were imaged and measured at pec-fin, day 5, and day 7 stages and p-values were calculated using the Welch two-sample t-test.

### Pigment quantification analysis in ImageJ

To assess variation of pigmentation across conditions, we used ImageJ(56) to measure the total area of pigmentation in larvae within the prim 20-25 stages and at the pec-fin stages, across treatment conditions. For this, lateral and dorsal view images were used to assess the level of pigmentation in epidermal melanocytes and in the RPE, respectively. Briefly, we employed a previously published pipeline(57) with minor modifications in the selection of the region of interest (ROI). We first converted all images to 16-bit images before selecting the ROI. We then created a mask from the ROI and followed by using the threshold function with default thresholding method. Next, we restored the selection and used the analyze particle function to measure the area of pigmentation within the ROI. For analyzing lateral view images, the contour of the larvae, without eyes and the yolk sac, was selected as the ROI. The eyes were excluded due to the superimposition of one eye over the other from a lateral view increasing the total pigmentation in one eye. The yolk sac, but not the yolk sac extension, was also excluded from lateral view analysis due to the level of shade in the periphery of the sac. The dorsal view images were utilized to analyze the total area of pigmentation in the eyes. Larvae were imaged and measured at prim 20-25 and pec-fin stages and p-values were calculated using the Welch two-sample t-test.

### T7 Endonuclease I (T7EI) assay for heteroduplex detection

Zebrafish embryos (n > 250) were injected with zebrafish-optimized *Cas9* mRNA (300 ng/µl, final concentration) that was *in-vitro* transcribed from the linearized pT3TS-nCas9n (Addgene #46757) plasmid vector. The microinjection mixture also included synthesized sgRNA mRNA (150 ng/µl, final concentration), nuclease-free water, and phenol red. At 48 hours post injection, clutches of 50 embryos per condition were collected for DNA extraction. DNA extraction was performed by incubating embryo clutches in 50 µl of zebrafish lysis buffer (1M Tris pH 8.4, 1M KCl, 1M MgCl_2_, 20% Tween-20, and 10% NP-40) at 94°C for 20 mins, 55°C (Hold to add 10 mg/ml Proteinase K), 55°C for 60 mins, 94°C for 20 mins, and optionally incubating at 4°C. Following extraction, DNA was amplified following standard PCR amplification protocols and purified using the Zymo Research DNA Clean & Concentrator-5 protocol (Cat. # D4013). The standard protocol from the T7EI assay from NEB (Cat. #M0689) was followed and after digestion, results were visualized in a 2% agarose gel.

### Ethics Statement

All zebrafish maintenance, handling, and analyses were performed under approved protocols from the Johns Hopkins University Animal Care and Use Committee (Protocol # FI22M203)

## AUTHOR CONTRIBUTIONS

A.S.M., V.C.C, and N.B.B. conceptualized the study. N.B.B. designed codon optimized dCas9-KRAB-MeCP2 and dCas9-VP64 constructs and custom single guide RNAs. N.B.B., E.A.S., and R.J.B. injected RNA constructs into zebrafish embryos. N.B.B. and E.A.S. imaged the fish. N.B.B. and R.J.B. wrote the manuscript. All authors edited and approved the final manuscript for publication.

## ACKNOWLEDGEMENTS

The authors would like to acknowledge the FInZ Zebrafish Core Center for assistance with zebrafish husbandry.

---

**Supplemental Figure 1:**
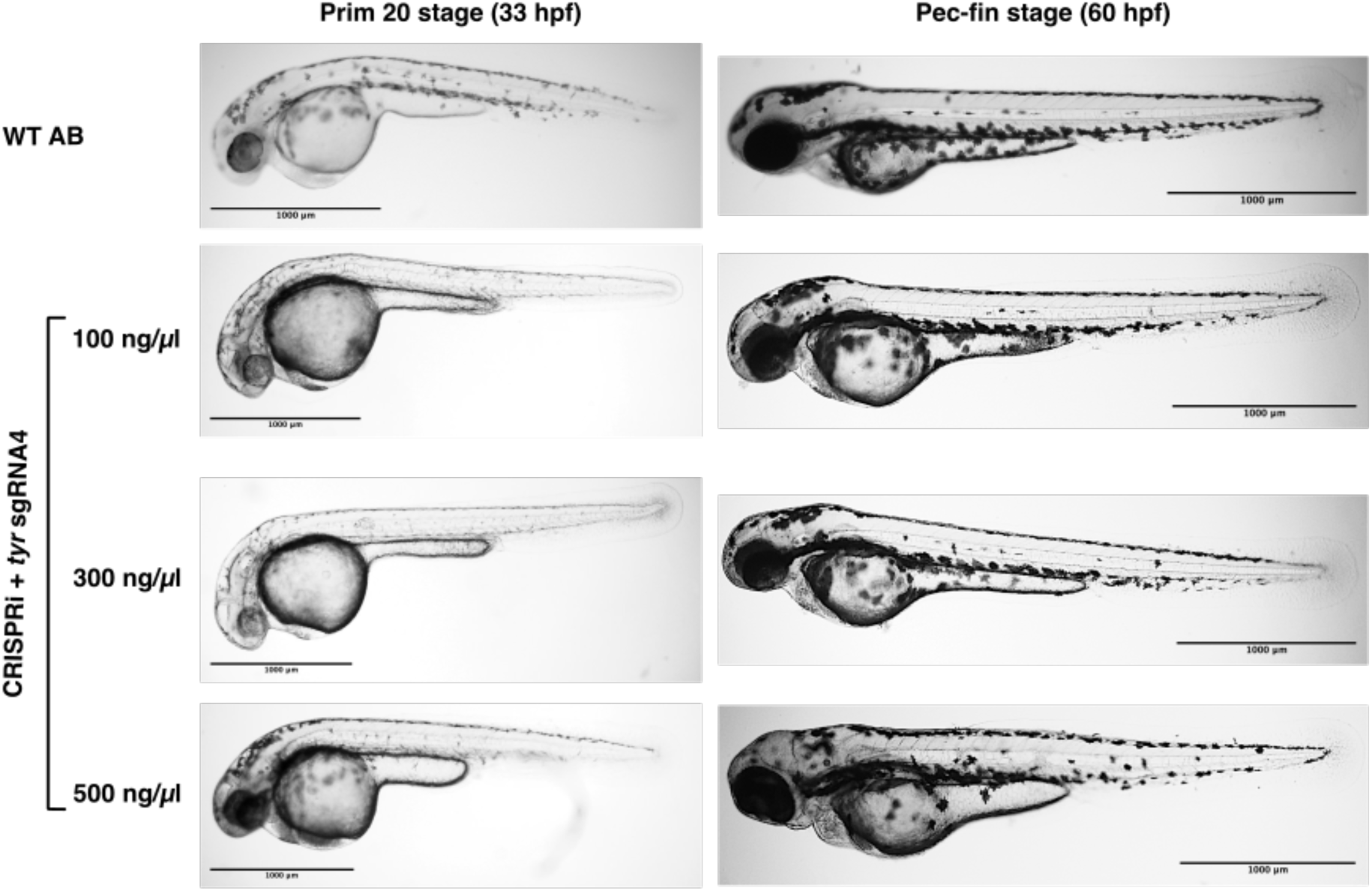
Assessment of optimal mRNA concentration for injection

**Supplemental Figure 2:**
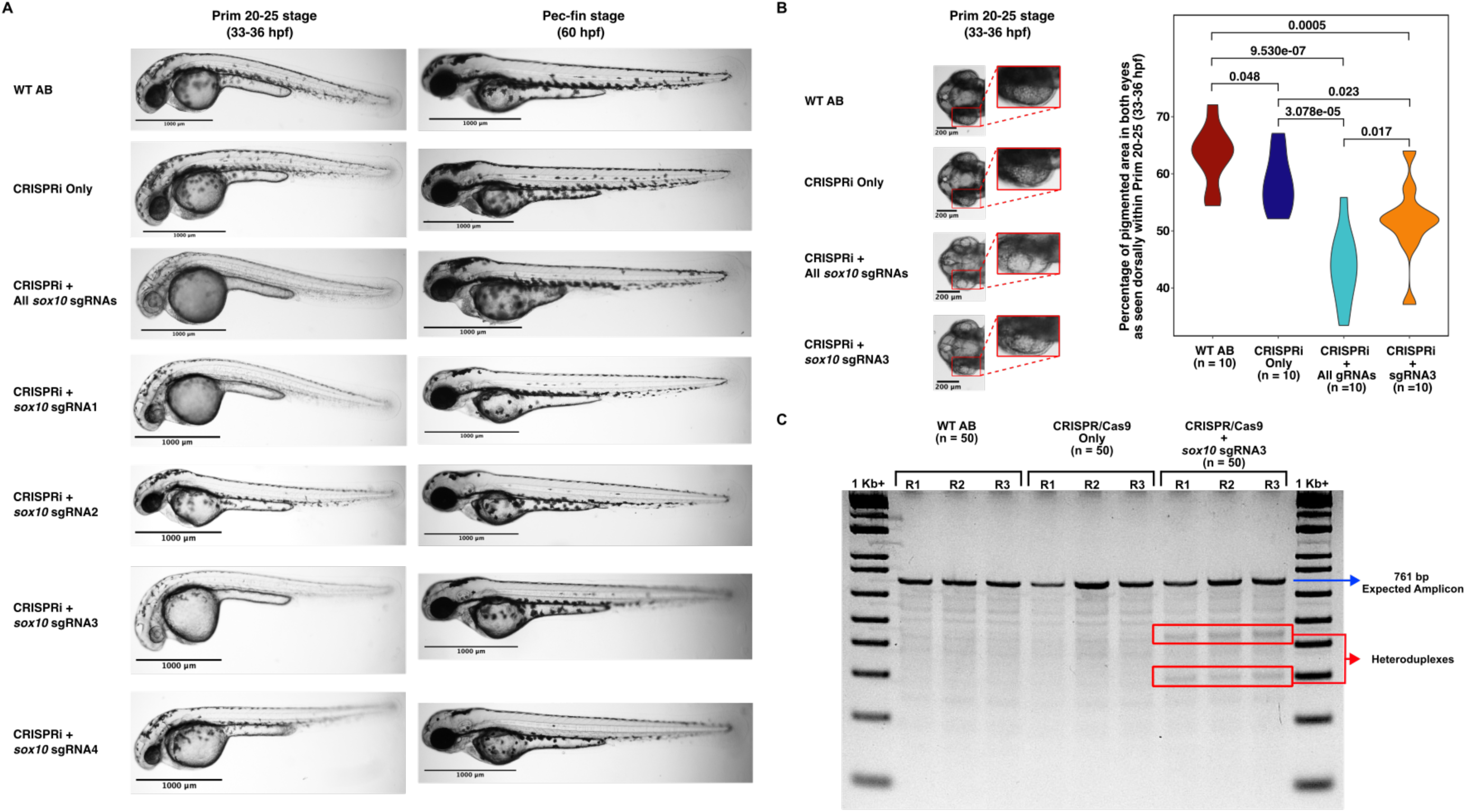
Targeting the *sox10* promoter with CRISPRi. **A.** Larvae at prim 20-25 (33-36 hpf) and pec-fin (60 hpf) stages across conditions. Scale bars 1000 µm. **B.** At the prim 20-25 stage, RPE pigmentation is variable across conditions; however, differences are statistically significant between controls and injected conditions. **C.** Gel (2% agarose) electrophoresis of T7 endonuclease I assay to detect heteroduplex formation (red boxes) after targeting the *sox10* promoter with sgRNA3. Expected amplicon length is 761 bp (blue arrow).

**Supplemental Figure 3:**
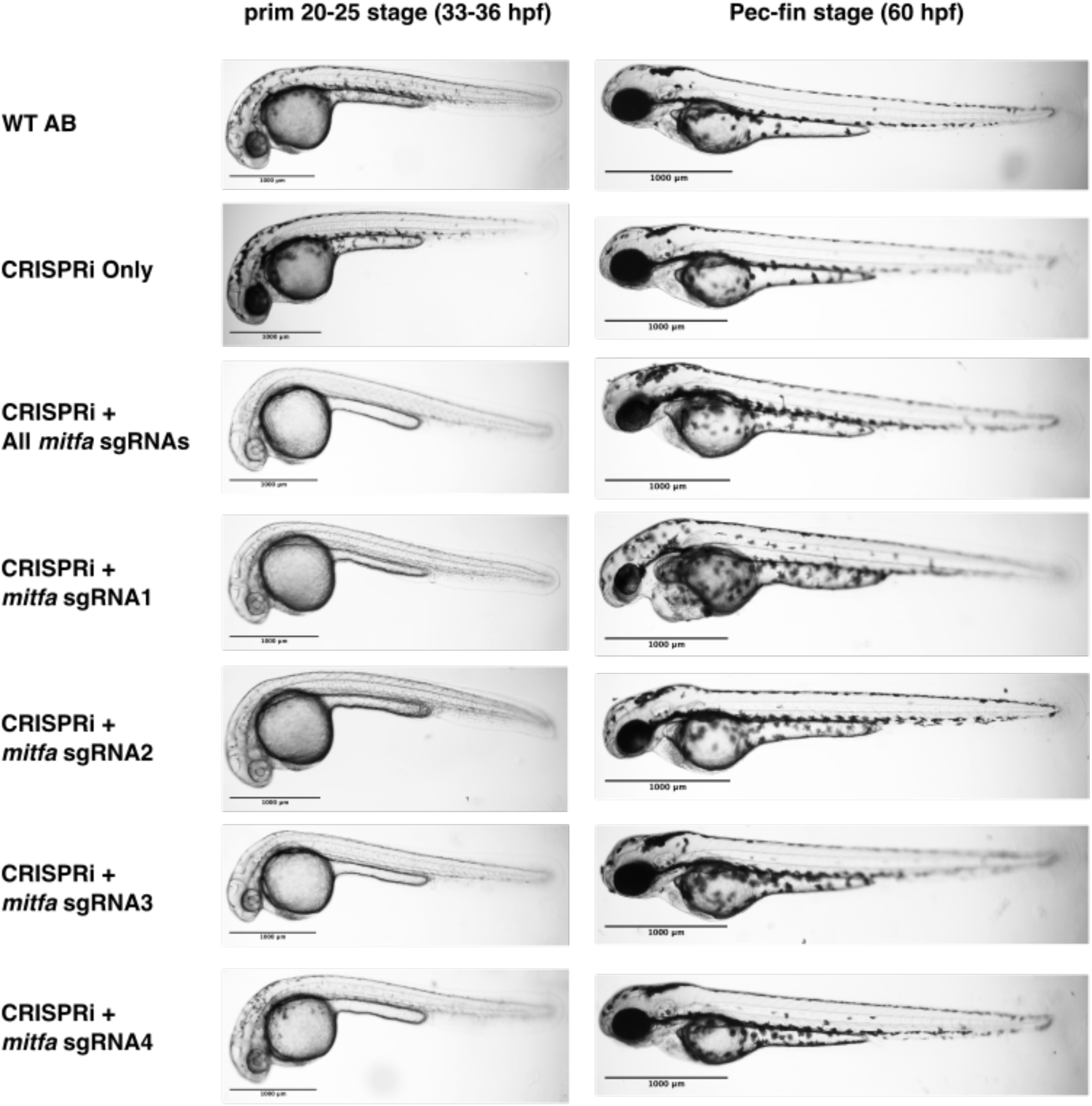
Targeting the *mitfa* promoter with CRISPRi. Scale bars 1000 µm.

**Supplemental Figure 4:**
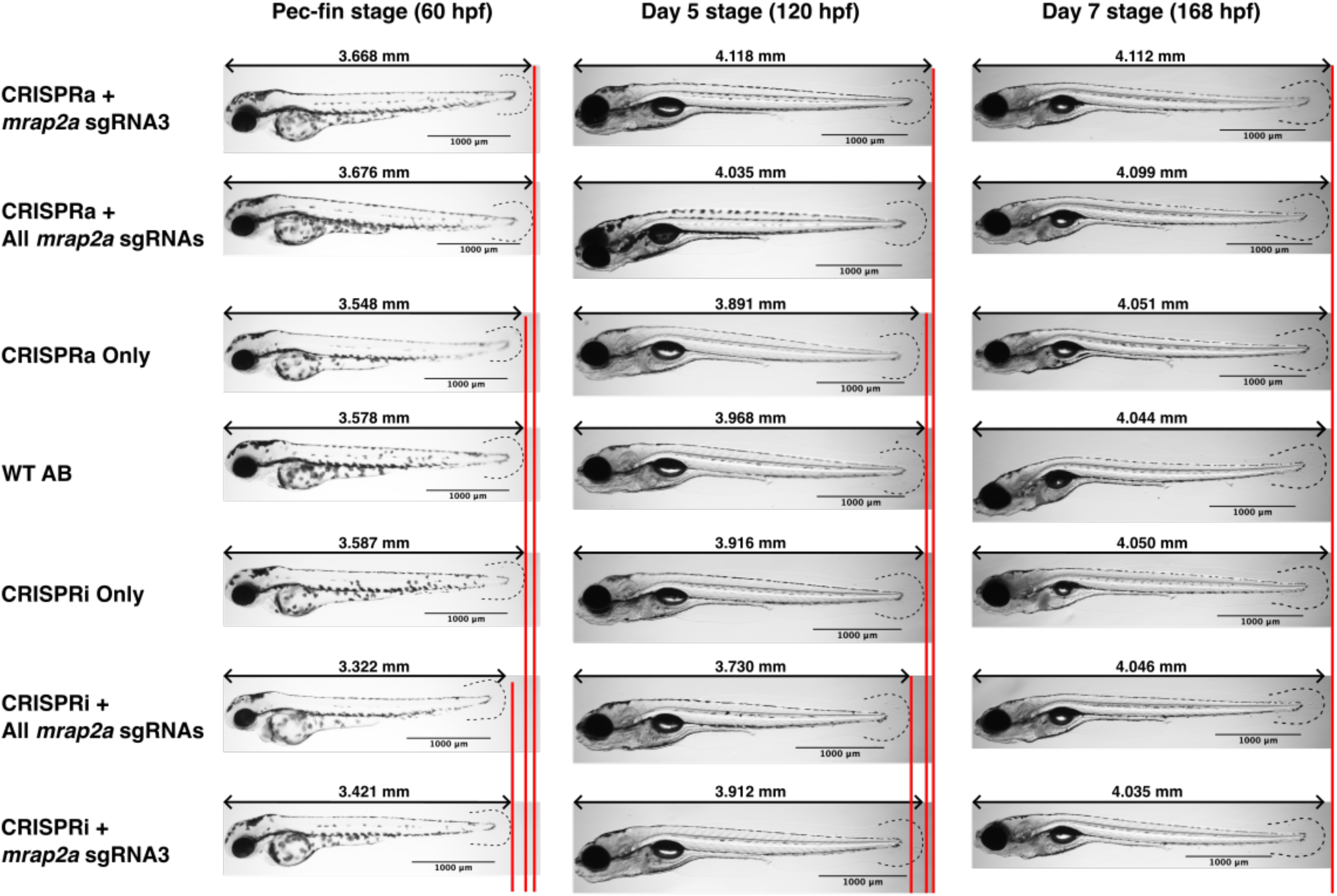
CRISPRa and CRISPRi targeting of the mrap2a promoter. Images taken at pec-fin, day 5, and day 7 stages. Dotted lines represent the translucent tail fin of the larvae. Scale bars 1000 µm.

